# AI-based identification of grain cultivars via non-target mass spectrometry

**DOI:** 10.1101/2020.05.07.082065

**Authors:** Kapil Nichani, Steffen Uhlig, Bertrand Colson, Karina Hettwer, Kirsten Simon, Josephine Bönick, Carsten Uhlig, Harshadrai M. Rawel, Manfred Stoyke, Petra Gowik, Gerd Huschek

## Abstract

Detection of food fraud and geographical traceability of ingredients is a continually sought goal for government institutions, producers, and consumers. Herein we explore the use of non-target high-resolution mass spectrometry approaches and demonstrate its utility through a particularly challenging case study – to distinguish wheat and spelt cultivars. By employing a data-independent acquisition (DIA) approach for sample measurement, the spectra are of considerable size and complexity. We utilize artificial intelligence (AI) algorithms (artificial neural networks) to evaluate the extensive proteomic footprint of several wheat and spelt cultivars. The AI model thus obtained is used to classify newer varieties of spelt, processed flour, and bread samples. Additionally, we discuss the validation of such a method coupling DIA and AI approaches. The novel framework for method validation enables calculation of precision parameters for facile comparison of the discriminatory power of the method and in the development of a reliable decision rule.

## Combating food fraud via DIA and AI

Public awareness of food fraud is mainly driven by high-visibility media discussions, e.g., in connection with public health consequences or when a large-scale operation is uncovered, and the ensuing scandal brings disrepute to companies or regulatory authorities. However, even when not topical, food fraud is widespread and exacts considerable economic costs. Its manifold manifestations include adulteration, mislabeling, dilution, substitution, etc. Establishing mechanisms and quality indicators to detect and limit the scope of food fraud, therefore, remains an important and urgent task.

Non-target mass spectrometry has been recognized as one of the most promising approaches for the purpose of identifying food grains and millets (Black et al. 2016). While many different non-target approaches exist, in the present paper, the focus will lie on data-independent acquisition (DIA) mass spectrometry. This is mainly due to the fact that DIA makes it possible to record a large ensemble of fragment ion spectra (Schilling et al. 2017). Spectra obtained via DIA are of considerable size and complexity and are best evaluated via artificial intelligence (AI) algorithms. For this reason, the methodology which is proposed here consists of a coupling of non-target approaches and AI data evaluation (artificial neural networks), with the aim of food fraud detection. In addition, the validation of such couplings is discussed, along with their modular nature.

## Benefitting from artificial intelligence

In DIA approaches, data are generated without any profiling constraints with respect to predefined peptides of interest. Accordingly, in order to unleash the full power of DIA approaches, intelligent processing of the time-resolved and mass segmented fragment ion maps is required. An efficient deconvolution of the manifold spectral features is achieved by applying AI methods to parse the complex and extensive spectra obtained from MS/MS fragments. Combining an AI-based approach with non-target high-resolution mass spectrometry (HRMS) measurements, we propose a method for fast, robust screening of grains for the identification of malpractice regarding labeling and nutritional diagnostics. In order to demonstrate the usefulness of such a method, we focus on two challenging grain types, which are genetically and morphologically similar – namely: spelt and wheat.

## Spelt vs. wheat

Spelts are derived from hybridization events between non-free threshing tetraploid wheat (2n = 4x = 28, genome = AABB) and free-threshing hexaploid wheat. They are very resilient to austere irrigation conditions. They have favorable digestive and nutritional values, driving up demand, and catapulting their market price. As these grains command a premium price, there is an economic benefit to devise methods for adulteration, tampering, substitution, etc. The general European Union (EU) legal framework, as put forward in regulations such as 2017/625 and 1169/2011, aims to ensure food safety and consumer protection by compelling producers to correctly label ingredients and their sources. In this case, product labelling must be combined with an authentication analysis of grain ingredients and additives. Declaration of authentication of wheat and spelt ingredients is a non-trivial problem, to say the least. Active efforts directed towards providing a framework to promote spelt varieties is only limited to a handful of countries. For example, in Germany there is a guideline (Leitsätze des Deutschen Lebensmittelbuchs für Brot und Kleingebäck) which serves as a guiding principle for the manufacture and sale of spelt bread. It states that spelt bread must contain at least 90% spelt. Similarly, in Switzerland guidelines laid out by IP-SUISSE and Bio-Suisse in cooperation with IG Dinkel regulate the growing and selling of spelt products. One particular challenge is the lack of a consensus in the EU regarding the definition of “spelt” and the scope of cultivars encompassing it. Furthermore, previously reported grain and cultivar type classification procedures are not fully satisfactory and can be applied to a selected number of cultivars only. Collectively, these aspects make authentication of products made from spelt varieties a formidable task, which only increases in complexity with the inclusion of “pre-spelt” and “wheat spelt” in processed foods.

## Non-target mass spectrometry

So far, it has been possible to achieve success in distinguishing individual spelt cultivars from wheat cultivars by using the protein patterns of Osborne fractions, gliadins, and glutens (detected by 2-D electrophoresis). Approaches based on MS fingerprinting should allow for broader applicability. While MS approaches often rely on the detection and quantification of specific peaks, discriminatory power can be improved by mobilizing the complete fragment-ion MS/MS spectrum rather than focusing on intensity values corresponding to a group of individual compounds.

For the analysis of the wheat, spelt, and crossed cultivars, protein extraction was followed by chymotrypsin digestion. These were measured using a UHPLC-Triple TOF (MS/MS) consisting of a micro-flow UHPLC expert microLC 200 with an autosampler CTC Pal system and a SCIEX ESI-TripleTOF 5600 with SWATH (sequential window acquisition of all theoretical fragment-ion spectra) acquisition. ProteinPilot was used for labeling the peptide sequences (complete experimental proteomics workflow as reported previously by Bönick et al., 2017 and Huschek et al., 2018). Although SWATH acquisition technology from SCIEX is employed here, the coupling of DIA and AI proposed is intended to include several other modes of DIA acquisition.

## From spectra to D-scores

The higher mass accuracy of SWATH-based acquisition requires elaborate data processing. A proprietary AI method was applied to leverage all the spectral information available via the non-target acquisition. Using a neural network-based approach, the algorithm allows the identification of cultivars based on the proteomic pattern. The neural network model learns by applying inductive reasoning to the correlations between different spectral features and corresponding structural features. Due to the sheer number of peaks involved, such correlations would be difficult for humans to recognize, let alone for inferences to be drawn in connection with a given identification objective. In the case of the identification of wheat vs. spelt, the AI method was trained on the basis of original spelt cultivars, i.e. that they are not crossed with wheat. The choice of original spelt cultivars, which have been bred decades ago, in training the AI model was motivated by the fact that different varieties have been cross-bred over the years to address specific issues in cultivation and processing, leading to highly mixed varieties. Hence using newer varieties would increase the difficulty involved in identifying the characteristic parts and features of the proteomic fingerprints. Each training sample was given an a priori identification value −1 (“spelt”) or 1 (“wheat”), which will take the role of a “true value” (See Figure 1). Each sample was “measured in duplicate”, meaning that two sets of spectra are available for each sample. For each spectrum, the proprietary AI algorithm then yields a standardized numerical value, called a D-score. The computation of these scores takes advantage of deep convolution nets for end-to-end learning of spectral features. For the proposed processing, parallels can be drawn to the wide-spread application of ANNs to image classification tasks. Higher-level abstractions of the input peak data through several convolutional and non-linear transforms are distilled to provide the D-score. The identification of a spectrum as “wheat” or “spelt” is then performed on the basis of the D-score.

**Figure 1:**
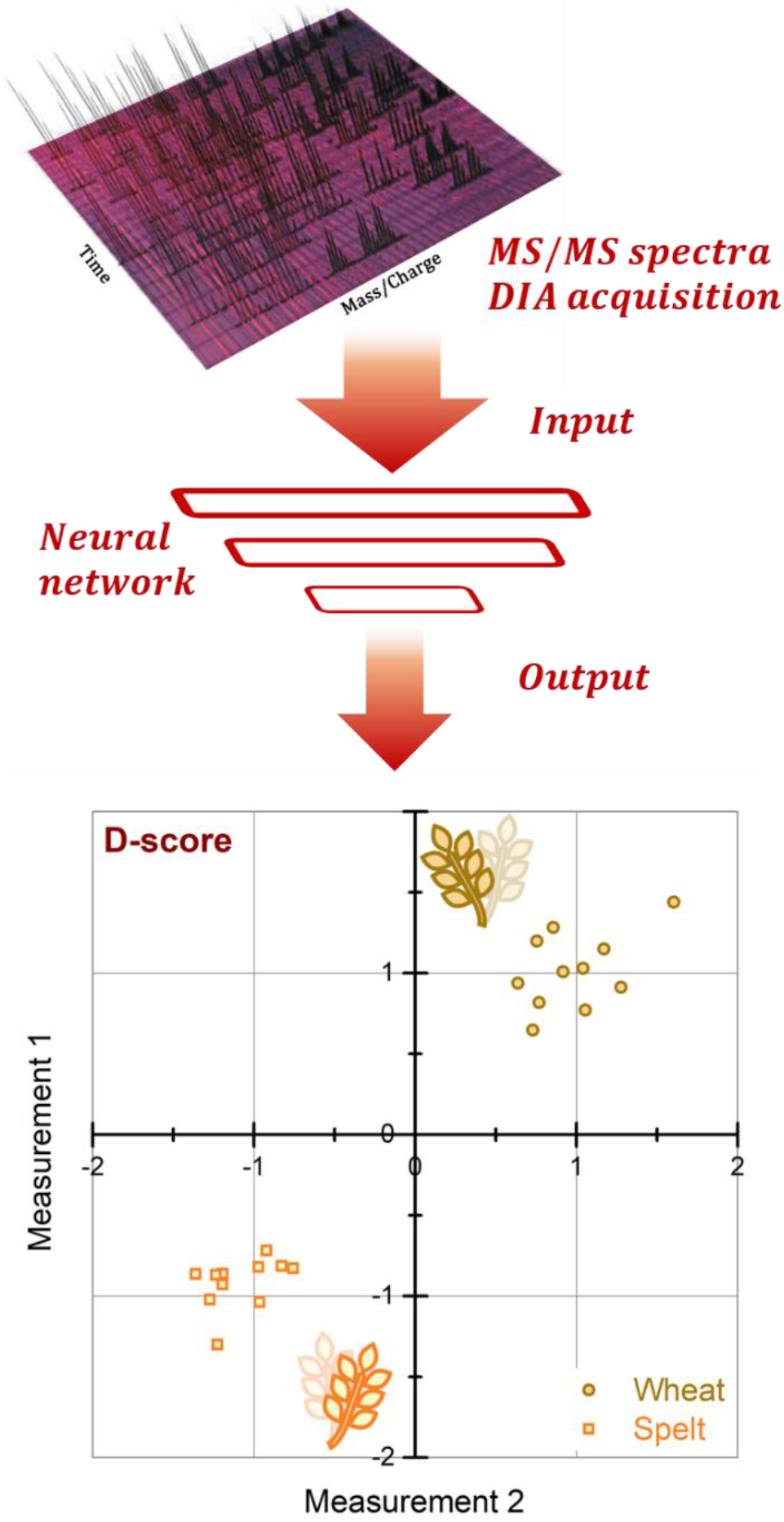
Schematic of the proposed method consisting of coupling DIA and AI stages. DIA spectra serve as an input to a neural network model. The bottom panel shows calculated D-scores for the cross-validation samples of 11 typical wheat and spelt cultivars, measured in duplicate, which are together used to train the model.

The validation of the AI method is performed simultaneously with model training using a specially adapted application of nested cross-validation. Often, relatively few samples are available for model training. Accordingly, it was essential to develop a new framework for the validation of the method and the evaluation of its performance characteristics. The trained model is further put to the test using different challenging real-world samples. Figure 2 shows a Youden plot with D-scores obtained for “atypical spelt” cultivars (comprising recently crossed varieties of spelt), processed flour mixture (in the form of a “spelt bread”) and “spelt flour”. Both of these samples contain spelt flour with 10% wheat (as per the guidelines set in Leitsätze des Deutschen Lebensmittelbuchs für Brot und Kleingebäck). Interestingly, we see heterogeneity in the D-scores of “old wheat” cultivars, bearing a closer resemblance to old spelt. In summary, the D scores for the tested cultivars demonstrate the functionality of our rapid, direct method for the identification of “spelt” vs. “wheat” varieties.

**Figure 2:**
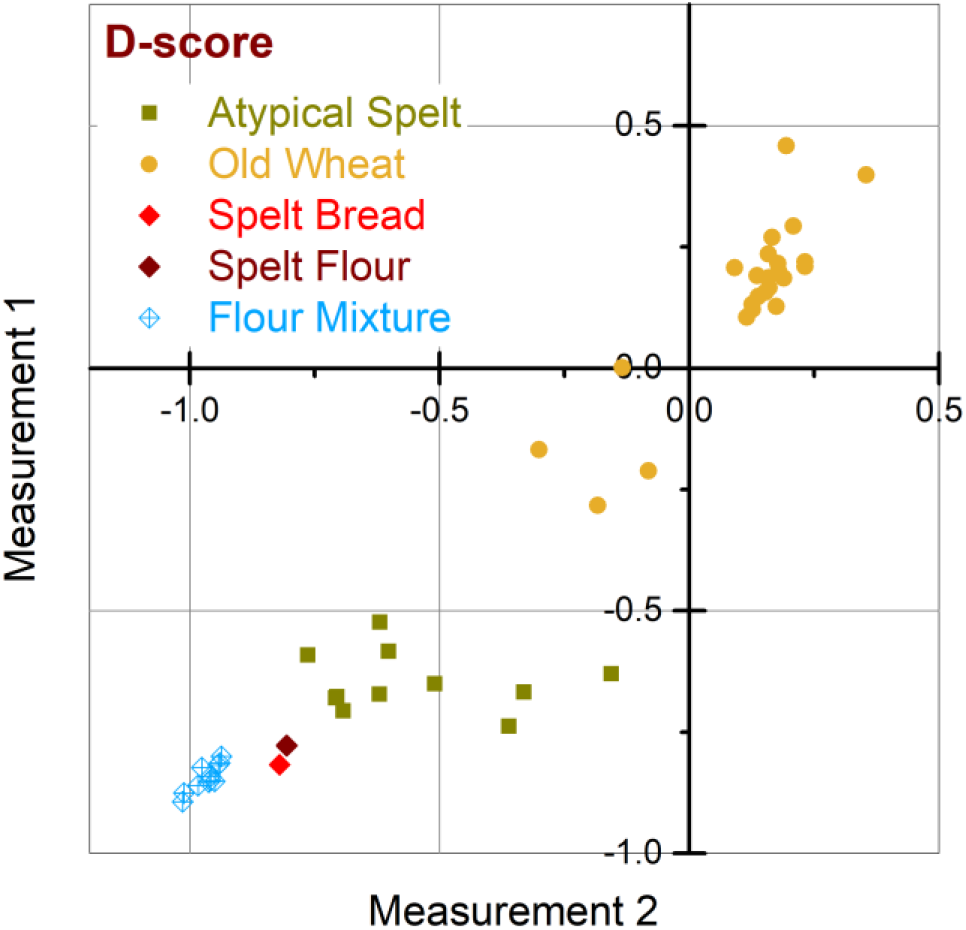
Youden plot with D-scores for the two “duplicate spectra” for some challenging samples to assess the applicability of the AI approach. Each point corresponds to one sample. A value close to 1 indicates that the method has identified the sample as wheat, whereas a value close to −1 indicates the sample was identified as spelt. Atypical spelt correspond to newer, cross-bred varieties. Old wheat collectively refers to traditional cultivars of wheat (around the year 1900). Spelt flour is 10% wheat flour in spelt and spelt bread is a baked product made with this spelt flour. Flour mixture samples were obtained by mixing spelt flour with wheat flour (10%) made from different grain types.

## Ensuring reliable decisions from scores

Method validation is a challenging problem for non-target mass spectrometry-based workflows. We propose a tech-nology-agnostic approach for the comprehensive validation of the method, understood as the coupling of the DIA and AI stages described above. It is essential to ensure that the discriminatory power remains adequate (a) when applied to other sets of data than the training set, (b) for the entire population falling under the scope of the method and (c) under all in-house testing conditions or when applied to data from different laboratories.

On the basis of the D-scores, different precision estimates can be obtained. Two of these precision estimates – called the population SD and (single laboratory) classification SD allow question (a) and (b) to be addressed. Other precision estimates correspond to more classical precision parameters such as repeatability, intermediate, and reproducibility SD, which allow aspect (c) to be addressed. The precision estimates for the D score can be obtained by using the approach described in previous reports (Uhlig et al. 2019 and Uhlig et al. Eurachem Workshop 2019). The single laboratory classification SD is used to check whether the decision rule can be considered reliable for the whole population falling within the scope of the classification method. SD values of 0.26 and 0.20 are obtained for “wheat” and “spelt” respectively. Since both values are well below 0.5, this suggests that the risk of misclassification is very low (<1%). With the intermediate SD, the reproducibility of the D score can be checked under in-house conditions: We obtained an intermediate SD of 0.18 for both wheat and spelt, which means that the analytical variability is almost equal to the variability between different wheat varieties. In contrast, the variability between spelt varieties seems to be much smaller. It can, therefore, be stated that the analytical variability is more than sufficient for the purpose of classification between wheat and spelt; on the other hand, the differences within the spelt cultivars studied are very small and cannot be precisely measured with the D score. The new approach is particularly beneficial in the case of low sample numbers, where more classical performance characteristics such as sensitivity, specificity, and ROC-AUC (area under the curve of the receiver operating characteristics, i.e., the curve which reflects the relation between sensitivity and specificity for varying thresholds) values could be less reliable. However, the selection of the cultivars studied was not strictly based on a random criterion. Due to the difficult availability of some varieties, the evaluation was done only on the basis of those cultivars that could be easily supplied. Therefore, the precision parameters can only be considered provisional. A confirmatory validation is still pending.

Another critical aspect is the availability of samples whose “true values” are known, as required for training. Knowledge about the real ID of the cultivar often relies on biochemical tests or known cross breeding history. Misconstrued labels will very likely result in reduced “generalizability” of the method.

## Conclusions and outlook

The methodology proposed in this paper consists of coupling DIA (e.g., SWATH) with AI data evaluation in order to obtain D-scores for different samples. These scores can then be used to tackle various questions in the fight against food fraud. The method can be easily adapted to allow the determination of geographical origin for grain samples, and this line of investigation is currently being explored. While the modular nature of the method can allow different stages to be replaced by alternative approaches (e.g., SWATH replaced by another DIA technology), the method validation principles would still be applicable, as stated here.

We aim to engage with instrument manufacturers and users to explore emerging application areas further. We see promise in the method’s usefulness not only in connection with the question of the authenticity of different food items and matrices but also in characterizing blood plasma in connection with diagnostic, prognostic, and therapeutic research.

